# The occurrence of *Aerococcus urinaeequi* and non-aureus Staphylococci in raw milk negatively correlates with *Escherichia coli* clinical mastitis

**DOI:** 10.1101/2024.03.12.584643

**Authors:** Dongyun Jung, Soyoun Park, Daryna Kurban, Simon Dufour, Jennifer Ronholm

## Abstract

*Escherichia coli* is a common environmental pathogen associated with clinical mastitis (CM) in dairy cattle. There is an interest in optimizing the udder microbiome to increase the resistance of dairy cattle to *E. coli* CM; however, the details of which members of the healthy udder microbiota may play a role in antagonizing *E. coli* are unknown. In this study, we characterized the bacterial community composition in raw milk collected from quarters of lactating Holstein dairy cows that developed *E. coli* CM during lactation, including milk from both healthy and diseased quarters (n=1,172). The milk microbiome from infected quarters was compared before, during, and after CM. A combination of 16S rRNA gene amplicon and shotgun metagenomic sequencing were used generate datasets with a high level of both depth and breadth. The microbial diversity present in raw milk significantly decreased in quarters experiencing *E. coli* CM, indicating that *E. coli* displaces other members of the microbiome. However, the diversity recovered very rapidly after infection. Two genera: *Staphylococcus, Aerococcus,* and the family *Oscillospiraceae* were significantly more abundant in healthy quarters with low inflammation. Species of these genera: *Staphylococcus auricularis, Staphylococcus haemolyticus,* and *Aerocussus urinaeequi* were identified by shotgun metagenomics. Thus, these species are of interest for optimizing the microbiome to discourage *E. coli* colonization without triggering inflammation.

**Importance:** In this study we show that *E. coli* outcompetes and displaces several members of the udder microbiome during CM, but that microbial diversity recovers post-infection. In milk from quarters which remained healthy, the community composition was often highly dominated by *S. auricularis, S. haemolyticus, A. urinaeequi,* and *S. marcescens* without corresponding increases in somatic cell count (SCC). Community dominance by these organisms, without inflammation, could indicate that these species could be potential prophylactics that could contribute to colonization resistance for the pathogen and prevent future instances of *E. coli* CM.

## Introduction

Bovine mastitis is an inflammation of the mammary gland of dairy cattle, mostly caused by intramammary infection (IMI). Mastitis is one of the most common and costly diseases in the dairy industry – it results in decreased quality and quantity of milk, expensive antibiotic-based treatments, and culling of chronically infected cows (1–3). Bovine mastitis causes an annual economic loss of $665 million CAD in Canada and $2 billion USD in the United States (2, 3). It is an extremely complex disease; 178 different bacterial species have been isolated from clinical mastitis (CM) samples and scientific literature describing their impact on somatic cell count (SCC) or clinical symptoms was available for 85 of these bacterial species (4). Different etiological organisms have different mechanisms of transmission and result in different symptoms. Based on modes of transmission mastitis causing organisms can be divided into two major categories: contagious or environmental; and based on symptomology mastitis can be clinical or subclinical (5). *Escherichia coli* is mainly associated with CM and was the most prevalent (11.0%) cause of CM in dairy cattle from Canada, US and Brazil, but can also be retrieved from apparently normal milking cows (i.e., subclinical mastitis) (4). It is primarily driven by environmental spread via feces, bedding, and soil on dairy farms (6). Despite improvements in hygiene practices on dairy farms, mainly aimed at controlling the spread of contagious pathogens, the number of mastitis cases caused by *E. coli* continues to increase annually (6).

*E. coli* mastitis is characterized by very acute, but generally transient infections of short duration that usually self-resolve without treatment (7–9) – although, persistent and recurrent mastitis caused by *E. coli* can occur (10). Since most cases of *E. coli* mastitis self-resolve, antibiotic treatment is not recommended (11, 12). However, severe cases can lead to septicemia and endotoxin-induced shock, and, therefore, an aggressive supportive therapy with, for instance, non-steroidal anti-inflammatory, fluids, and, possibly, systemic antibiotic, is important (9, 13, 14). Moreover, *E. coli* isolated from cattle with bovine mastitis have been shown to be resistant to several classes of antibiotics including aminopenicillins, polypeptides, lincosamides, and macrolides (15–18). Therefore, intramammary administration of third-generation cephalosporins, specifically ceftiofur, is commonly used by producers to treat cases of severe *E. coli* CM, even though the added-value of such a treatment is not demonstrated (12, 14, 19, 20). However, extended-spectrum β-lactamase (ESBL) and plasmid-mediated AmpC β-lactamase-producing *E. coli* have now been reported in cattle with mastitis, raising concerns about both human and bovine health (21–24).

Significant effort has been put into developing vaccines targeting *E. coli* mastitis in dairy cattle, and vaccination specifically against coliform mastitis has been a part of control programs for three decades (25). However, there has been limited uptake of the available vaccines in commercial herds (26), antibody levels are known to decline over time (27), and, to maintain their efficacy, most of these vaccines have to be administered repeatedly at relatively short time-interval, which limits their practical application.

Probiotics may be an effective way to prevent bacterial infections in the bovine mammary gland (28–30). Lactic acid bacteria (LAB) have been tested as probiotic candidates and inhibited colonization by bovine mastitis pathogens and inflammatory responses *in vitro* and *in vivo*. *Lactococcus garvieae, Lactococcus lactis, Lactobacillus brevis, Lactobacillus casei, Lactobacillus platarum* and *Lactobacillus perolens* inhibited colonization by *E. coli, Staphylococcus aureus,* and *Streptococcus uberis* and inflammatory responses of bovine mammary epithelial cells induced by *E. coli in vitro* (28–31)*. L. lactis* was also able to inhibit the growth of *S. aureus in vivo* (32). However, despite the effectiveness *L. lactis in vivo* it is unknown if *L. lactis* is maintained as part of the microbiota long enough to protect the host indefinitely (32). More importantly, LAB in milk can be responsible for organoleptic defects in cheese and other dairy products (33) and therefore, care should be taken when developing LAB probiotics for the dairy industry.

Optimizing the udder microbiome has been suggested as a promising way to protect dairy cattle from mastitis (34, 35), since the microbial population that inhabits mammals as part of a healthy microbiota provides a layer of protection against pathogen colonization as well as the overgrowth of opportunistic pathobionts (36). The bovine udder is home to a rich microbial community; although, the composition of the bovine udder microbiome varies based on the site (37–40). The teat apex, teat canal, raw milk, and raw colostrum each have distinct microbiomes and a healthy udder microbiome tends toward a high level of diversity, while a mastitic quarter tends toward a single or a small group of bacterial species (35, 39–41). The milk microbiome from a healthy cow generally contains *Firmicutes*, *Proteobacteria*, *Bacteroidetes*, and *Actinobacteria* at the phyla level, and *Staphylococcus*, *Ruminococcaceae*, *Lachnospiraceae*, *Propinoibacterium*, *Stenotrophomonas*, *Corynebacterium*, *Pseudomonas*, *Streptococcus*, *Comamonas*, *Bacteriodes*, *Enterococcus*, *Lactobacillus*, and *Fusobacterium* at the genus level (42–46). Certain members of the udder bacterial community may reduce susceptibility to mastitis via one of several mechanisms. Some non-aureus staphylococci (NAS), such as *Staphylococcus chromogenes*, have been shown to antagonize growth of *Staphylococcus aureus* (47, 48). The protective effects of NAS likely occur due to the ability to produce bacteriocins (49, 50), purine analogues (51), or biofilm disrupting signals (52). The growth of Gram-negative mastitis pathogens has not been shown to be inhibited by NAS (48). Certain NAS are also known to cause mastitis – which complicates their potential use as probiotics (53). Compared to work with *S. aureus*, relatively little work has been conducted to identify bovine commensals which may provide protection against *E. coli* CM. *Bacillus subtilis* has been shown to limit damage to mammary tissues during *E. coli* CM – although, this protective effect appears to be caused by immune modulation as opposed to inter-bacterial antagonism (54).

In this study, we hypothesized that the raw milk microbiome in dairy cattle quarters that remained healthy would have clear and consistent differences compared to the milk from quarters that developed *E. coli* CM, and that, in identifying these differences, we may identify members of the udder microbiome that are specifically antagonistic toward *E. coli*. To test this hypothesis, we collected bi-weekly quarter-level milk samples from a cohort of dairy cattle that were also concurrently being monitored for *E. coli* CM. We characterized the longitudinal changes in the composition of the bacterial community in raw milk samples and performed a differential analysis comparing quarters that experienced *E. coli* CM during the study to quarters that remained mastitis-free using 16S rRNA gene-targeted amplicon sequence (TAS) analysis. To further identify members of the microbiome which negatively correlated to *E. coli* CM and further understand the *E. coli* strains that were causing CM we selected certain milk samples to analyze via shotgun metagenomics. Overall, we determined that the presence of *Staphylococcus, Aerococcus*, and UCG-005 each had a negative correlation to the occurrence of *E. coli* CM.

## Results

### Sample information, sequencing coverage, and overall bacterial community composition

In this study, 19.3% (135/698) of cattle developed CM during the study period. Of the 135 diagnosed CM infections, 19.25% (26/135) were caused by *E. coli*. Mastitis cases caused by *E. coli* are described in Fig. 1 and Table S1. The first *E. coli* CM cases were reported across all stages of lactation: four during the transition period (1-21 days of milk (DIM)), ten in early-lactation (22-100 DIM), seven in mid-lactation (101-200 DIM), and five in late-lactation (>201 DIM). A total of 1,336 quarter-level milk samples were collected from the 26 cows diagnosed with *E. coli* CM (Table S2; Fig. 1) and DNA extraction and 16S rRNA sequencing was performed on each sample. During data processing 209 samples were removed from the study due to low Good’s coverage (< 99.0%) or a low number of sequences reads (<3,100) (Table S3). Therefore, 1,127 milk samples were included in the analysis alongside kit controls (n=15) and negative PCR controls (n=15). In addition, 41/1,127 samples produced more than two different colony morphologies on blood agar. Based on guidance of the National Mastitis Council milk samples which produce more than two colony morphologies on blood agar should not be analyzed due to possible contamination (Table S4) (55). We chose to sequence these samples despite possible contamination and analyzed data twice both including and excluding these 41 samples. Results obtained from data including these 41 samples are reported throughout and were, for the most part, identical to results excluding these samples; however, it if results differed between the two analysis it is indicated. Sequencing efforts produced total of 3,497,437 sequence reads passed filter (Table S3). Contamination in the negative control was not identified (Fig. S1).

**Fig 1.**
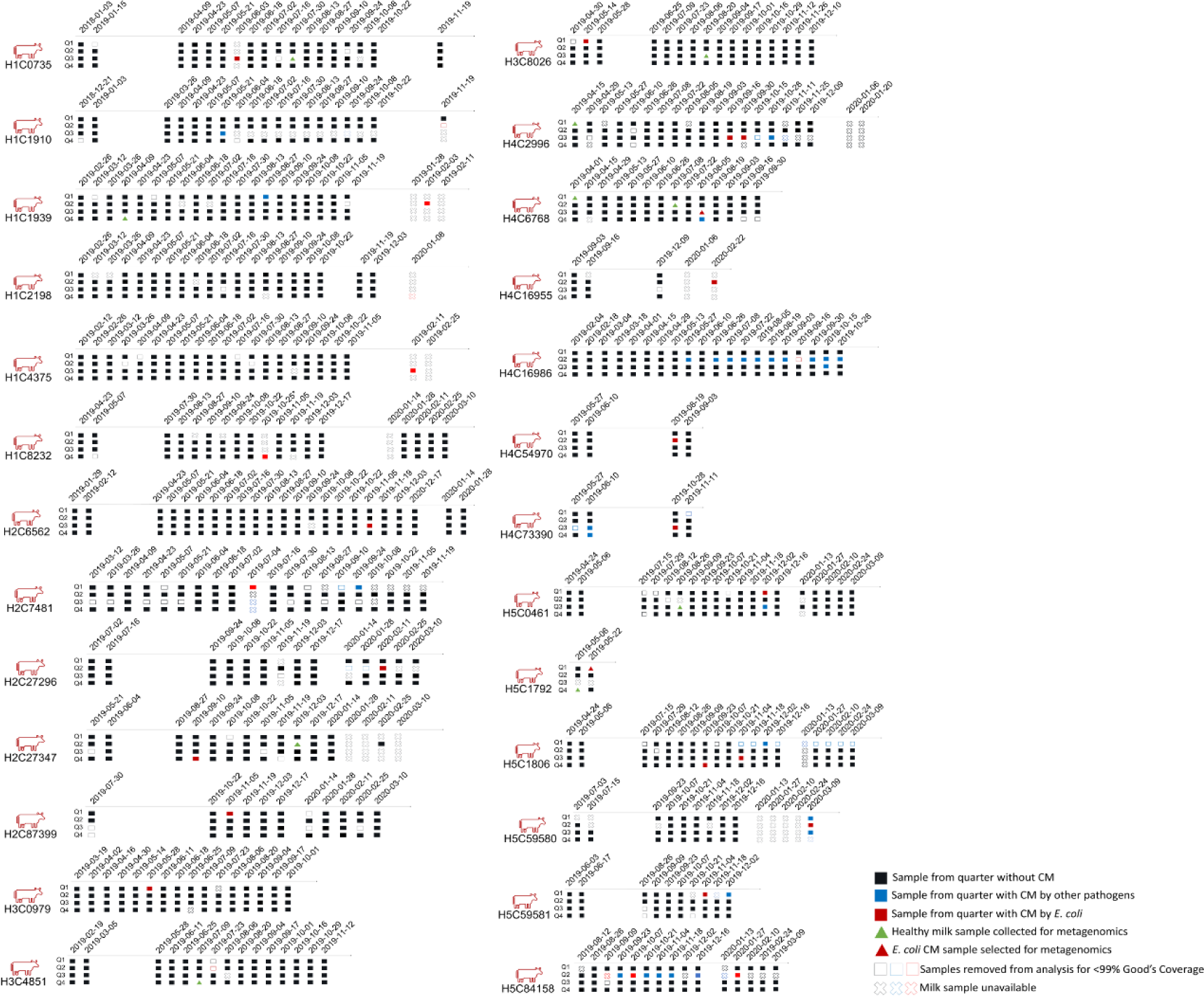
Twenty-six Holstein cows with *E. coli* clinical mastitis (CM). Our study followed 26 Holstein cows that developed *E. coli* CM during the study from five herds in Quebec, Canada. Raw milk samples were taken from each quarter of the cows bi-weekly. The H and C on each cow ID indicate the herd and assigned cow number (ATQ). A total of 1,172 milk samples were included in the microbiome analysis; the red and blue boxes represent the samples from the quarters that had CM caused by *E. coli* and other mastitis pathogens, respectively. The black boxes represent the samples from the quarters without CM. The samples indicated with empty boxes were excluded from the analysis due to low Good’s coverage (< 99.0%) or low sequence reads (< 3,100). The samples indicated with X were unavailable for DNA extraction due to insufficient volume (> 1.0 mL).

Milk samples from five herds shared four main phyla: Firmicutes, Proteobacteria, Bacteroidota, and Actinobacteriota (Fig. 2A and 2B). Firmicutes were the most dominant group across all the samples with an average relative abundance of 51.45% followed by Proteobacteria (23.73%), Bacteroidota (11.46%), and Actinobacteriota (9.80%) (Fig. 2A). At the genus level, five most abundant OTUs represented *Staphylococcus, Aerococcus, Escherichia*_*Shigella,* UCG-005 from the Oscillospiraceae family, and an unclassified genus from Enterobacteriaceae family (Fig. 2C). Chao1 index was significantly different between herd 1 and herd 5, and herd 3 and herd 5 (TukeyHSD post-hoc test; *p* < 0.01) (Fig. S2). The permutational multivariate analysis of variance (PERMANOVA) analysis of the Bray-Curtis dissimilarity showed that beta diversity was significantly different between herds (PERMANOVA; *p* < 0.01) (Fig. S2).

**Fig 2.**
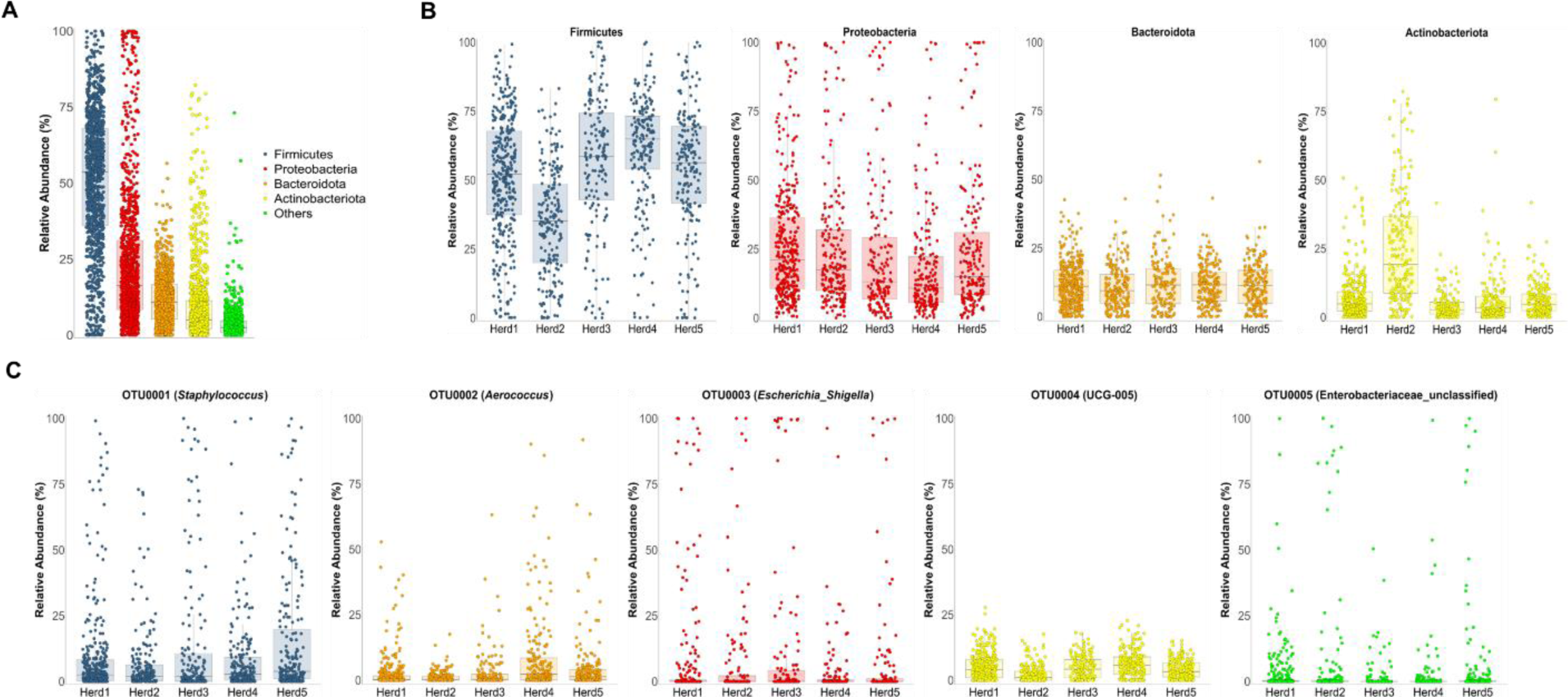
Relative abundance of the raw milk microbiota from 26 Holstein dairy cows that developed *E. coli* CM. (A) Firmicutes, Proteobacteria, Bacteroidota, and Actinobacteriota were the major phyla identified in the milk samples. (B) The relative abundance of each of the four phyla varied between the five herds, except Bacteriodota which was consistent between herds. *Staphylococcus* was the most abundant OTU, followed by OTUs associated with *Aerococcus, Escherichia_Shigella,* UCG-005, and Enterobacteriaceae_unclassified genera.

### Microbial changes in milk before, during, and after *E. coli* CM

The diversity of the milk microbiome was compared over time between quarters which developed *E. coli* CM and quarters that remained free from *E. coli* CM. Alpha-diversity was not significantly different between these categories of quarters if the milk was collected two-weeks before or after *E. coli* CM; however, alpha-diversity was significantly different on the day of *E. coli* CM diagnosis (Shannon *p* < 0.01; Mann–Whitney statistic 84.5, Simpson *p* < 0.01; Mann-Whitney statistic 86.5, Chao1 *p*<0.01; Mann-Whitney statistic 196.5) (Fig. 3A and Table 1). However, when samples were analyzed without the 41 samples that produced more than 2 colonies on blood agar, Chao1 was not significantly different on the day of *E. coli* CM (Chao1 *p*=0.0104; Mann-Whitney statistic 236.0). The difference in alpha-diversity on the day of CM was driven primarily by the dominance of the *Escherichia-Shigella* OTU (Fig. S3 and Table 1). Differences in beta-diversity were also only significant on the day of CM based on PERMANOVA analysis of the Bray–Curtis dissimilarities (PERMANOVA *p* < 0.01, F = 15.169) (Fig. 3B). The relative abundances of OTU0001 (*Staphylococcus*), OTU0002 (*Aerococcus*), and OTU0004 (UCG_005) were significantly higher in milk taken from healthy quarters compared to milk taken from *E. coli* CM quarters on the day *E. coli* CM was diagnosed (Mann-Whitney U test adjusted by Benjamini & Hochberg (BH); *p* < 0.01) (Fig. 3C; Fig. S3). These correlations were also identified using linear discriminant analysis (LDA) where *Escherichia_Shigella* was highly associated with milk from quarters with *E. coli* CM (LDA=5.49), and *Staphylococcus*, *Aerococcus*, and UCG-005 were highly associated with milk from healthy quarters (LDA < −4.1) (Kruskal-Wallis rank-sum test; *p* < 0.01) (Fig. 3D). Other OTUs were also associated milk from healthy quarters on the day of *E. coli* CM, including: *Ruminococcaceae*, *Bifidobacterium*, *Bacteroides*, *Atopostipes*, and *Jeotgalicoccus* (LDA > 2.0 or < −2.0; KW rank-sum test *p* < 0.01) (Table 1).

**Fig 3.**
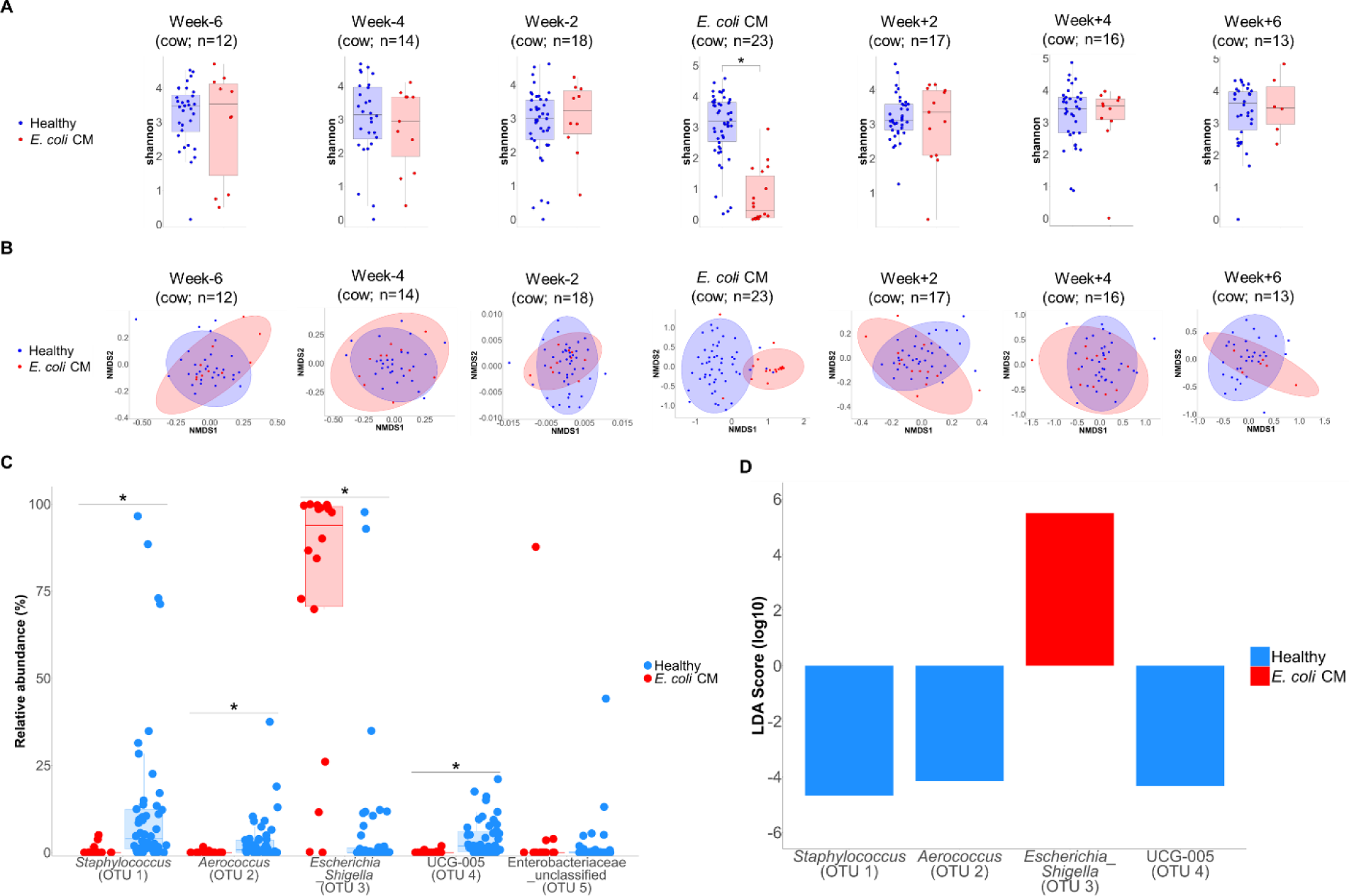
Microbial community changes in alpha- and beta-diversity in milk collected before, during, and after the first *E. coli* CM case. In (A) and (B) both the alpha- and beta-diversity of the microbial community in milk are similar in healthy and *E. coli* CM quarters before and after the disease. However, significant differences were demonstrated on the day of CM in milk from the infected quarter ((A) Shannon *p* < 0.01 and Mann–Whitney statistic 84.5, (B) Permutational multivariate analysis of variance (PERMANOVA) *p* < 0.01, F = 15.169)). In (C) and (D) differential abundance analysis was conducted on the relative abundance of the five most abundant OTUs from the first *E. coli* CM using Mann-Whitney U test and LEfSe analysis. The analysis identified that OTU0001 (*Staphylococcus*), OTU0002 (*Aerococcus*), OTU0004 (UCG-005) were significantly associated with healthy quarters; and OTU0003 (*Escherichia*_*Shigella*) was significantly associated with *E. coli* CM quarters.

**Table 1.**
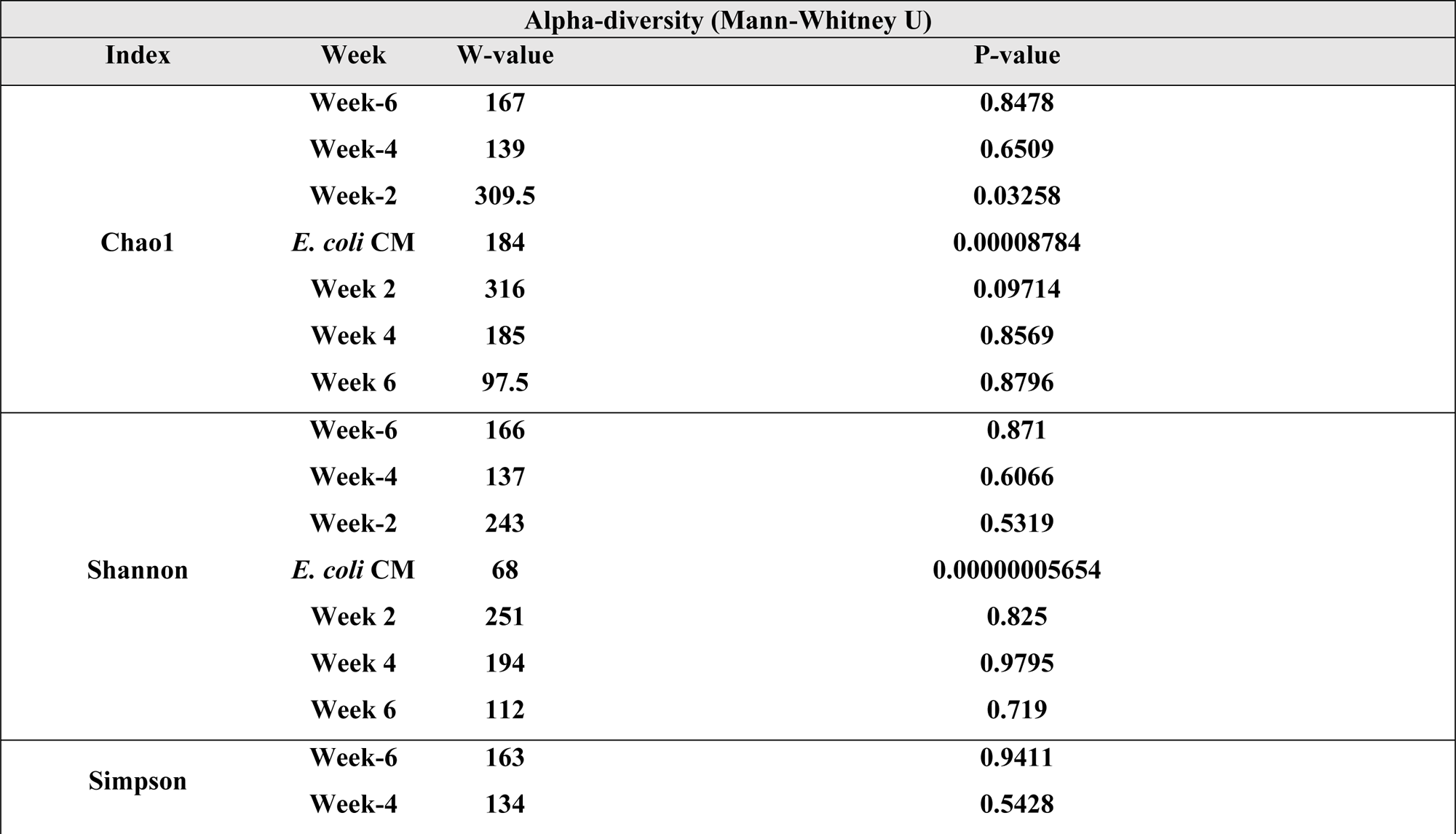
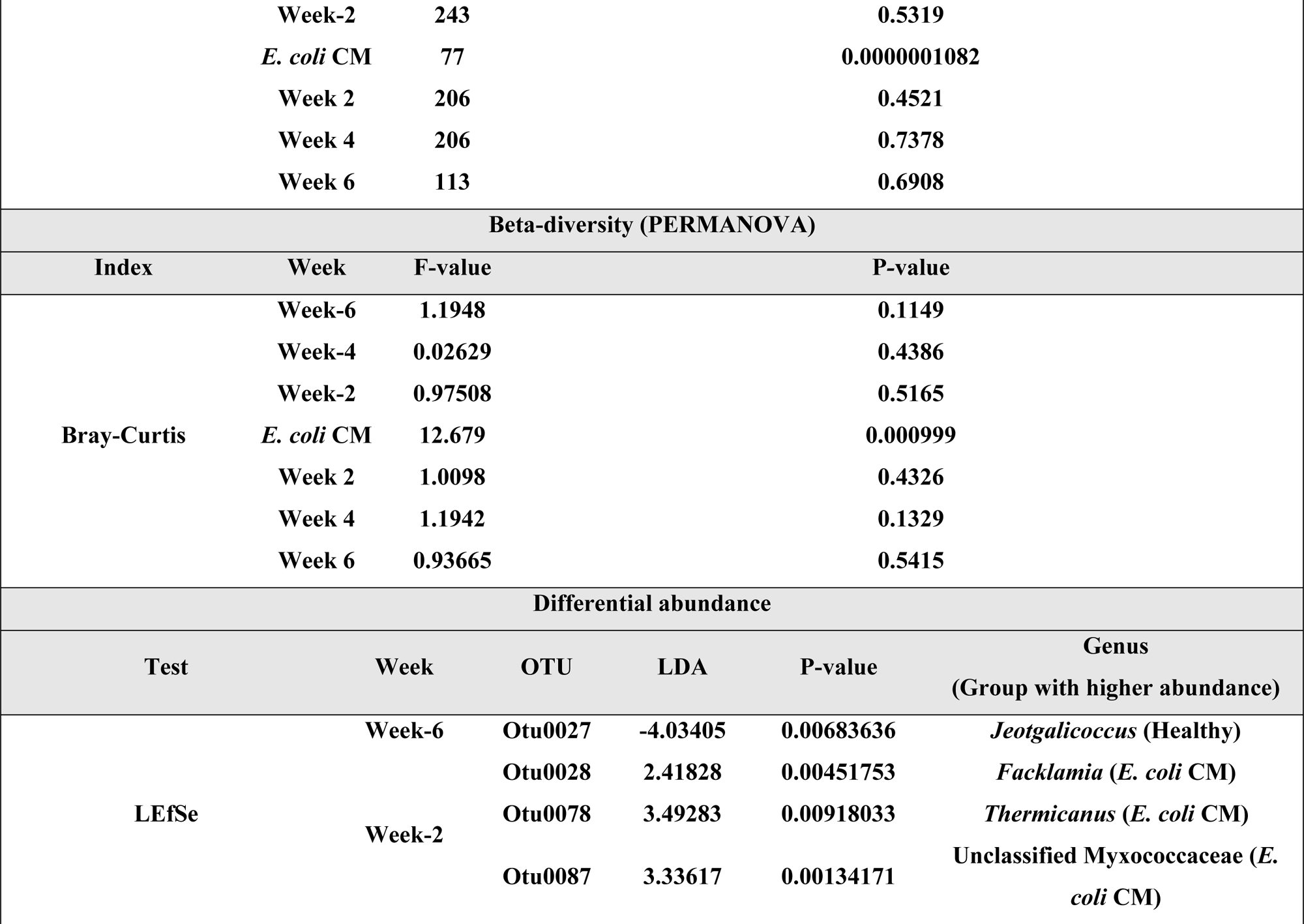
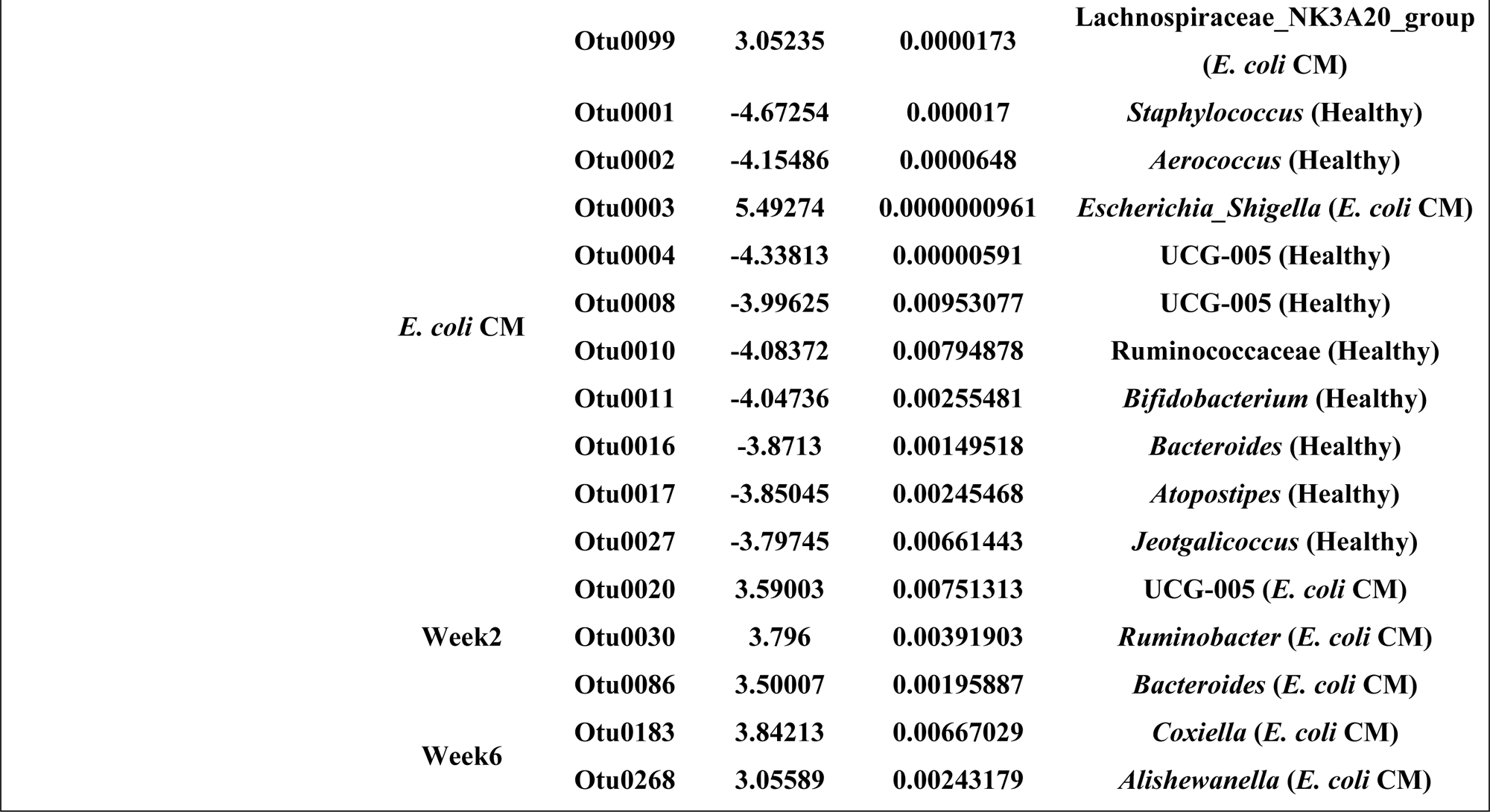
Statistical analysis on microbial changes in milk microbiota before, during, and after the first *E. coli* clinical mastitis (CM) case.

### Microbial network analysis of the bacterial community

To determine potentially biologically relevant interactions that occur between each of the OTUs found in milk, the 187 OTUs which had more than 10% of prevalence across 1,127 samples were further investigated (Fig. S4 and Table S5). No significant associations were identified between any OTUs (−0.165 < Spearman’s ρ < 0.666).

### Correlations between SCC and bacterial abundance and diversity

The correlation between the log SCC and the Shannon index in milk samples from quarters that did not have *E. coli* CM (n=844) was evaluated to determine if there was a clear relationship between bacterial alpha-diversity and inflammation in the absence of CM. The Shannon index of the samples had a weak correlation with the log SCC (Spearman’s ρ = 0.009) (Fig. 4A). The correlations between the log SCC and each of the five most abundant OTUs in healthy samples were also evaluated, but all OTUs showed a weak correlation (0 < Spearman’s ρ < 0.1) (Fig. 4B)

**Fig 4.**
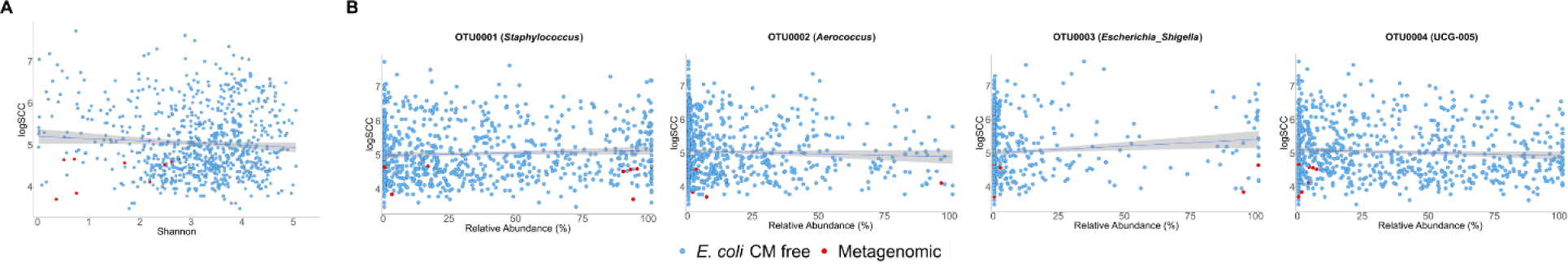
Correlation between SCC and the raw milk microbiota. (A) Shannon index was not significantly correlated with log SCC (Spearman’s ρ = 0.009). (B) The relative abundance of OTU0001 (*Staphylococcus*) (Spearman’s ρ = 0.099), OTU0002 (*Aerococcus*) (Spearman’s ρ = 0.023), OTU0003 (*Escherichia*_*Shigella*) (Spearman’s ρ = 0.072), and OTU0004 (UCG-005) (Spearman’s ρ = 0.033) also had only weak correlation with log (SCC). There were mastitis-free milk samples that had more than 40% relative abundance of these OTUs and less than 50,000 cells/mL SCC (log(SCC)=4.7). Some of these milk samples were selected for shotgun metagenomic sequencing, and samples selected for shotgun metagenomics are shown in red.

### Shotgun metagenomic analysis of milk from healthy quarters

Select samples (n=11) with a high relative abundance (> 40%) of each of the two most common OTUs and a low SCC (< 50,000 cells/mL; log SCC < 4.7) were selected for shotgun metagenomic sequencing to identify bacteria, at the species level, that may outcompete *E. coli* in this environment but do not result in inflammation (Fig. 4B). A total of 476,966,298 raw sequence reads were produced (Table S6). Filtering was preformed to remove any sequence reads that corresponded to bovine DNA. After bovine DNA was removed 36,259,190 sequence reads remained (Table S7). Three MAGs: *Aerococcus urinaeequi* (Genome size: 1,859,582 bp; CDS: 1,683; completeness = 97.80%, contamination = 1.10%), *Staphylococcus auricularis* (Genome size: 2,006,007 bp, CDS: 1,889; completeness = 98.34%, contamination = 0.0%) and *S. haemolyticus* (Genome size: 2,623,558 bp; coding sequences (CDS): 2,603; completeness = 98.48%, contamination = 1.99%), were assembled from sample 30000729. Sample 30000729 was taken from an exceptionally healthy quarter that did not develop CM from any pathogen during the study period and had an SCC of 5,000 on the day the sample was collected.

Overall, taxonomic profiling was performed on these 11 metagenomes identified the presence of 12 species including: *Acinetobacter gandensis, A. lwoffii, A. towneri, A. urinaeequi, C. bovis, C. stationis, Glutamicibacter arilaitensis, Luteimonas sp J29, P. fluorescens, Serratia marcescens, S. auricularis* and *S. haemolyticus* (Fig. 5 and Table S8).

**Fig 5.**
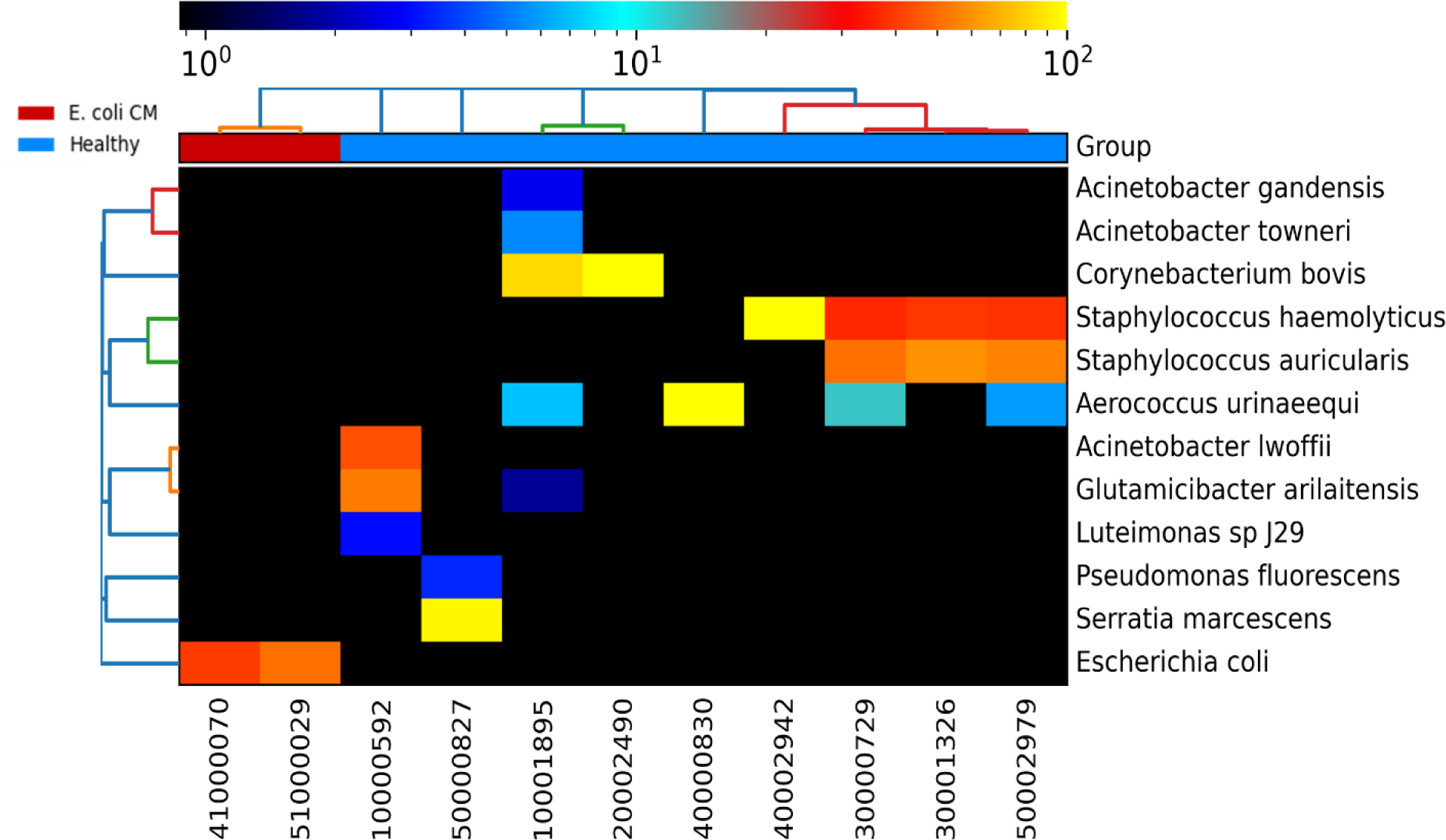
Microbial composition of healthy and *E. coli* CM samples at species-level. The relative abundance of taxonomic composition of the shotgun metagenomic samples was estimated by MetaPhlAn4 using the species-level genome bins (SGBs) database (mpa_vJan21_CHOCOPhlAnSGB_202103) with the default parameters. The abundance of 13 species higher than 0.1% of relative abundance was identified from 11 healthy and two *E. coli* CM samples.

Milk sample 50000827 was selected for metagenomic sequencing since it was identified as having a high abundance of the *Escherichia-Shigella* OTU (90.42%), but a low SCC (44,000). This led us to question if certain strains of *E. coli* could act as commensal organism in the bovine udder and not result in inflammation. There were several milk samples in our dataset that had a high relative abundance of *Escherichia-Shigella* but low SSC (Fig. 4). Metagenomic sequencing revealed a single MAG of *S. marcescens* (Genome size: 2,840,286 bp, CDS: 2,672; completeness = 71.29%, contamination = 1.84%) from this sample. To verify the similarity of V4 region in 16S rRNA gene between *E. coli* and *S. marcescens,* we aligned the sequences of the V4 region (Fig. S5). The alignment showed 98.8% identity, which was above the threshold to assign OTUs, indicating that *S. marcescens* is also included in this OTU. Therefore, no evidence of a commensal mammary *E. coli* was identified in this investigation.

Annotation of the MAGs identified several genes which could contribute to *E. coli* antagonism including the D-lactate dehydrogenase gene, *ldhA,* in the MAGs of *S. haemolyticus* and *S. marcescens*, and the L-lactate dehydrogenase gene, *ldh*, in the MAGs of *S. auricularis, S. haemolyticus*, and *A. urinaeequi*. The genes of class IIa bacteriocins listeriocin and pediocin were identified in MAG of *A. urinaeequi.* The *S. marcescens* MAG contained the genes of ten core components (TssA, TssE, TssF, TssG, TssH, TssJ, TssK, TssL, TssM) and two accessory components (Hcp, VgrG) of a type VI secretion system (T6SS).

### Shotgun metagenomic analysis of milk from *E. coli* CM Quarters

Two milk samples with *E. coli* CM (41000070 and 51000029) were selected for metagenomic sequencing to provide additional insights into the pathogen and the composition of the bacterial community during *E. coli* CM. A total of 113,859,360 raw sequence reads were produced from these two samples (Table S6). Filtering was preformed to remove any sequence reads that corresponded to bovine DNA. After bovine DNA was removed 172,994 sequence reads remained (Table S7). All sequence reads were identified as *E. coli*.

## Discussion

We investigated the bacterial composition in raw milk taken before, during, and after *E. coli* CM from both healthy and CM quarters of 26 dairy cows that developed natural *E. coli* CM during our study period. We conducted 16S rRNA amplicon sequencing to track the changes of microbiota in the milk samples from the quarters that developed *E. coli* CM and the quarters that remained healthy for the study. Shotgun metagenomic analysis was conducted on the samples taken from healthy quarters to profile potential commensal bacteria at the species level that may outcompete or be actively antagonistic toward *E. coli*.

The microbiota in raw milk samples was dominated by four phyla: Firmicutes, Proteobacteria, Bacteroidota, and Actinobacteriota. We observed the variations in the abundance of these taxa across herds and cows except Bacterioidota which remained at a constant low level. This observation is similar to previous studies that demonstrated microbiome variation from the farm-to-farm level to the cow level (56–58). Actinobacteriota was significantly more abundant in herd 2 than in the other herds. *Glutamicibacter*, a genus from Actinobacteriota, accounted for most of the Actinobacteria in herd 2. The number of *E. coli* CM cases in herd 2 was not significantly different from other herds. Some genera of Actinobacteriota phyla, such as *Streptomyces* and *Bifidobacterium*, have been reported to have antimicrobial activities against *E. coli* (59, 60), however, each was present at very low relative abundances in our dataset.

We investigated changes in bacterial community composition in the samples taken from all quarters of 26 cows before, during, and after the first *E. coli* CM, and observed that milk samples taken during active *E. coli* CM mastitis had significantly decreased alpha-diversity, but this quickly rebounded after the infection. This indicates that pathogen clearance and the reestablishment of the udder microbiome happen quickly after infections, which has been observed previously (43, 61). In earlier work, experimental infection with *E. coli* led to a peak in *E. coli* counts 16-24 hours post-infection, and bacterial clearance without treatment seven days after a challenge, although, about 50% of cows intermittently shed *E. coli* asymptomatically until the end of the study period (61). A decrease in bacterial diversity in milk samples prior to CM has been observed for other mastitis pathogens, and may even be predictive for a future infection in the case of *S. aureus*, but observations from our study indicate that diversity alone is not predictive of which quarters will develop *E. coli* CM (37, 56).

The differential abundance analysis of various bacterial taxa between healthy samples and samples taken from quarters with *E. coli* CM identified a significant increase in the abundance of *Escherichia-Shigella* in mastitic milk and a significant increase in the abundance of *Staphylococcus*, *Aerococcus*, and UCG-005 in healthy samples. *Staphylococcus*, *Aerococcus*, and UCG-005 were also highly dominant in some healthy samples that had less than < 50,000 cells/mL of SCC (Fig. 4B). The dominance of these genera in healthy quarters, without triggering inflammation, may imply these members may be able to outcompete pathogenic species in this environment and should be investigated further as potential probiotics. The lack of a significant association between the SCC and Shannon index and the high relative abundance (>40%) of some OTUs in the healthy samples that had less than 50,000 cells/mL of SCC also implies a small impact of dysbiosis on the susceptibility to the disease.

To find the specific species of *Staphylococcus*, *Aerococcus*, and UCG-005 which could dominate a microbial community without causing inflammation, we performed metagenomic sequencing of samples with a high abundance of *Staphylococcus* or *Aerococcus* but a low SCC. This identified *S. auricularis, S. haemolyticus,* and *A. urinaeequi* at the species level. *S. auricularis* and *S. haemolyticus* are NAS and have previously been reported to cause subclinical mastitis and CM (53, 62–68). However, a previous study reported that *S. auricularis* and *S. haemolyticus* could be present on the teat apices of dairy cows without causing IMI (62); other recent studies found that NAS makes up a significant portion of the healthy udder microbiota and that they are predominant in the raw milk samples (SCC < 200,000 cells/mL) from the healthy quarters of Holstein dairy cows (53, 56, 69). Furthermore, NAS species including *S. capitis, S. chromogenes, S. epidermidis, S. pasteuri, S. saprophyticus, S. sciuri, S. simulans, S. warneri,* and *S. xylosus* can inhibit the growth of *S. aureus* associated with clinical or subclinical mastitis by bacteriocins and the purine analog 6-thioguanine (49–51). However, the inhibition of growth of *E. coli* by NAS has not been previously reported (48).

The MAGs of *S. auricularis, S. haemolyticus,* and *A. urinaeequi* were constructed from a healthy milk sample, and each genome encodes D- and L-lactate dehydrogenase which catalyzes the production of lactic acid from pyruvate. Several studies have reported that lactic acid can inhibit the growth of *E. coli* O157:H7 *in vitro* (70–72). Considering the previous studies that tested LAB as probiotic candidates to prevent IMI and the high prevalence of L-lactate dehydrogenase across the healthy samples (28–32, 73), *S. auricularis, S. haemolyticus,* and *A. urinaeequi* could be capable of prevention while being maintained long enough as commensal groups.

*A. urinaeequi* has rarely been identified as a bovine mastitis pathogen while *Aerococcus viridans,* a close phylogenetically related species (74), is known as a subclinical mastitis pathogen (75, 76). *A. urinaeequi* has demonstrated antimicrobial activity against *Klebsiella pneumoniae, Salmonella enterica, Vibrio alginolyticus,* and *S. aureus* (56, 77). A newly identified class II bacteriocin from *A. urinaeequi* has shown antagonistic activity against *K. pneumoniae, S. enterica,* and *V. alginolyticus*, but the activity against *E. coli* by the bacteriocin has not been identified yet (77).

A MAG identified as *S. marcescens* was constructed from a healthy milk sample (5000827); the annotation of the MAG identified genes of T6SS components comprising the genes of ten core components (TssA, TssE, TssF, TssG, TssH, TssJ, TssK, TssL, and TssM) and two accessory components (VgrG and Hcp). Previous studies found that *S. marcescens* could have T6SS and inhibit the growth of *E. coli in vitro* (78, 79). *S. marcescens* is an etiological agent of bovine clinical mastitis (80). However, the milk sample we identified as containing a high abundance of *S. marcescens* had a SCC of 44,000 cells/mL, indicating a relatively healthy quarter. These findings could indicate that this strain of *S. marcescens* does not illicit significant inflammation.

In this study, we characterized changes of the microbial community before, during and after *E. coli* CM in the bovine udder and identified commensal bacteria in the samples from healthy quarters correlate negatively to the presence of *E. coli.* An unbalanced microbiota, driven by the overgrowth of certain commensals, is not necessarily associated with intramammary inflammation and bovine mastitis. In our study, we identified four such species, and the assembled MAGs of these species each encoded mechanisms that can potentially antagonize *E. coli*. The modulation of the microbiome with these species could be a potential way of preventing future *E. coli* CM without inducing inflammation. Further studies need to focus on the isolation of these species from the healthy cows from a wide range of geographical origins and their antagonistic effect on *E. coli in vitro* and *in vivo* as potential probiotics.

## Materials and Methods

### Sample collection

The study was affiliated with Park *et al.,* as part of a larger project that included 698 Holstein dairy cows from five herds in the Quebec province, in proximity to the Faculty of Veterinary Medicine at Université de Montréal in Saint-Hyacinthe (56). Raw milk samples were collected aseptically from each quarter every two weeks during the lactation period between December 2018 and March 2020. Briefly, the farm’s teat disinfectant was applied, after a contact time of 20 seconds, the teats were dried using paper towel and each teat end was then scrubbed with individual alcohol swabs. Approximately 60 mL of milk was then discarded, and 50 mL were collected (i.e., cisternal milk). Milk samples were immediately refrigerated at 4°C and transported, on the same day, to the Faculty of Veterinary Medicine laboratory where they were aliquoted for the different parts of the study (i.e., one aliquot for routine microbiological culture, one aliquot for microbiome analyses, and one aliquot for SCC and milk components measurements). In parallel to these farm visits, CM was identified by the producers by visual inspection of the milk and udders, and a milk sample was collected by the producers at diagnosis, following the protocol previously described. These latter samples were immediately frozen on the farm at −20°C, collected by the research team during the next farm visit, and processed in the laboratory as previously described.

For each milk sample, 10 µL per sample was spread on 5% sheep blood agar followed by 24-48 hours of incubation at 35°C, and number of phenotypically different CFU was noted (81). Milk sample with more than two phenotypically different CFU on blood agar were considered contaminated (55). The scores from 0 to 3 were assigned to no growth, one pure culture with one species, mixed culture with two species, and contaminated with three or more different species, respectively. The species of the colonies on the agar were identified except the ones with the score 3 without β-hemolysis (Table S4). Isolates were collected from the blood agar plates spread with CM samples to identify the etiological agents of mastitis. Bacterial species were identified using matrix-assisted laser desorption/ionization time-of-flight (MALDI-TOF) mass spectrometry. All the collected milk samples were then stored between −10°C and −20°C until bacterial DNA extraction.

### Bacterial DNA extraction

Bacterial DNA was extracted from each milk sample taken from an animal that developed *E. coli* CM during the study period. The samples were taken from six weeks before to six weeks after the first *E. coli* CM (Fig 1). The DNeasy^®^ PowerFood^®^ Microbial Kit (QIAGEN, Germany) was used following the manufacturer’s instructions with minor exceptions. To prepare milk samples for DNA extraction, frozen milk was fully thawed on ice, and inverted several times to mix, and then a 1 mL aliquot was placed in an Eppendorf microcentrifuge tube centrifuged at 16,000×g for 10 minutes. The supernatant was discarded, and the DNA extraction protocol was carried out using the pellet. Separate negative and positive extraction controls were included in the analysis for each reagent kit used. For a negative control, the extraction procedure was carried out using DNA/RNA-free water in place of a milk pellet. The positive control consisted of extracting DNA, using the same kit, from a rumen sample from a generous donor. To extract a higher quantity of DNA for shotgun metagenomic sequencing, a 6.0 mL aliquot of milk was used. The concentration and purity of extracted DNA from all samples and the controls were evaluated using Invitrogen™ Quant-iT™ dsDNA Assay Kit (Thermo Fisher Scientific, USA) and a Nanodrop 2000 (Thermo Scientific, USA), and samples with poor quality or quantity were re-extracted.

### 16S rRNA gene TAS

The V4 region of the 16S rRNA gene from the extracted bacterial DNA from milk samples (n=1,336) and controls (n=30) were amplified using PCR with the F515 and R806 primer pair and sequenced using Illumina MiSeq platform (Illumina Inc., San Diego, CA, USA) (82). The HotStartTaq® Plus Master Mix Kit (QIAGEN, Germany) was used for PCR and the amplification cycle included initial denaturation at 95°C for 5 minutes, 35 cycles of denaturation at 95°C for 30 seconds, annealing at 50°C for 30 seconds, extension at 72°C for 1 minute, and final extension at 72 °C for 10 minutes. The amplicon libraries were purified using Agencourt AMPure^®^ XP (Beckman Coulter, Brea, CA, USA) as per the manufacturer’s instructions and quantified using Invitrogen™ Quant-iT™ dsDNA Assay Kit. The libraries were then normalized and pooled (>1 nM) followed by sequencing using the MiSeq reagent kit V2 for 502 cycles (2 x 251bp) and the MiSeq benchtop sequencer.

### 16S rRNA gene TAS data analysis

FASTQ files were generated from the Illumina MiSeq sequencer and analyzed using Mothur (v. 1.42.3) (83). Mothur aligned the read pairs to make contigs with the length 253 bp and the similar contigs were clustered and aligned by the SILVA reference (v. 138) (84). Then, chimeric sequences were removed using UCHIME, and the filtered contigs were clustered and assigned to operational taxonomic units (OTUs) at 97% of identity as a threshold (85). The average OTUs were obtained by rarefying the sequences 1,000 times repeatedly to the minimum number of sequences (n=3,100) using Vegan R package (vegan::rarefy.perm) (v. 2.6-2) (86, 87). Following the rarefaction, Good’s coverage of the sequences was calculated on R (v. 4.1.2), and all samples with higher than 99.0% Good’s coverage were retained for analysis (88).

### Diversity and relative abundance of abundant taxonomic groups analysis

Differences in alpha-diversity (Chao1, Shannon, Simpson) and beta-diversity (Bray-Curtis) were compared between milk samples collected from healthy quarters and quarters that developed *E. coli* CM using the Vegan R package. The Mann-Whitney U test and PERMANOVA test was conducted to identify significant differences in alpha- and beta-diversities between the sample groups. Non-metric multidimensional scaling (NMDS) ordination was used to plot the beta-diversity using the Vegan R package. The Mann-Whitney U test and linear discriminant analysis effect size (LEfSe) were conducted to determine the differential abundance of OTUs between the two types of samples. The OTUs with LDA scores higher than 2.0 from LEfSe and BH adjusted *p-*value lower than 0.01 were considered to be significant (89). Spearman correlation and standard linear regression were used to characterize the relationships between log (SCC) and Shannon index, and between log (SCC) and the abundant OTUs on R.

### Microbial network analysis

The microbial network from all milk samples was created using the rarefied OTU table in R. Each network was created by calculation of co-occurrence using the Spearman correlation between the OTUs and corroborated with two OTU-generalized linear models (GLM) — one that included only environmental independent variables and another that included independent variables and the relative abundance of each other’s OTUs (56, 90). Cows and quarters as environmental independent variables and Quasipoisson distribution on each OTU-OTU combination were used for GLM analysis. The OTU-OTU correlations that had a *p-*value >0.01, discrepant results between the Spearman and GLM analyses, and a Spearman’s ρ ≤ 0.2 and ≥ −0.2 were excluded (90, 91). The standardized β coefficient values were used to visualize the networks on Cytoscape (v. 3.8.2) (92).

### Shotgun metagenomic sequencing

The samples selected for shotgun metagenomic sequencing included two milk samples taken from *E. coli* CM quarters on the day of diagnosis, and 11 milk samples from healthy quarters that had more than 40% relative abundance of one of the five most abundant OTUs and low SCC (< 50,000 cells/mL). After DNA extraction, the Nextera XT DNA Flex Library Preparation Kit (Illumina Inc., San Diego, CA, USA) and Nextera XT Index Kit were used to prepare metagenomic libraries following the manufacturer’s instructions. Then, paired-end sequencing (2 x 150 bp) was performed using the Illumina NovaSeq 6000 sequencer by Genome Quebec (Montreal, QC, Canada).

### Metagenomic analysis

BioBakery tools were used for the analysis of the metagenome sequences (93). Raw FASTQ files were processed using KneadData (v. 0.10.0) which filtered out Illumina Nextera adapters, and the host genome (https://github.com/biobakery/kneaddata). Bos taurus 3.1 (UMD 3.1, https://bovinegenome.elsiklab.missouri.edu/downloads/UMD_3.1) was a reference host genome. The threshold of the quality score was 30, and highly duplicated sequences (> 40%) were de-duplicated using the clumpify.sh function by BBMap (v.38.86) (94). The metagenomic data was analyzed using both genome binning and assembly-free approaches.

For genome binning, the filtered reads were assembled by metaSPAdes (v. 3.15.4) followed by taxonomic binning by metaBAT2 (v. 2.14) and MaxBIN2 (v. 2.2.7) (95–97). The bins were refined using MAGpurify (v. 2.1.2) and evaluated by CheckM (v. 1.0.18) which filtered out the bins with less than 70% of completeness and more than 2.0% of contamination (98, 99). The species of metagenome-assembled genomes (MAGs) from the successful bins were identified using the Genome Taxonomy Database Toolkit (GTDB-Tk) (v. 2.2.3) with the reference database, R207_v2, and the annotation was done using Prokka (v. 1.14.5), BAGEL4, AntiSMASH and RASTtk (v. 2.0) on BV-BRC (v. 3.30.19) (100–104).

For assembly-free analysis, the filtered reads after KneadData were processed by MetaPhlAn 4 (v. 4.0.3) to identify the taxonomic groups at the species-level using the species-level genome bins (SGBs) database (mpa_vJan21_CHOCOPhlAnSGB_202103) with the default parameters (105–107). Heatmaps of the taxonomic profiles were plotted using hclust2 with the default parameters except for Bray-Curtis as the distance function for samples (https://github.com/segatalab/hclust2).

## Accession number(s)

All 16S rRNA TAS and shotgun metagenomic sequences have been deposited and are available at the NCBI Sequence Read Archive under BioProject accession numbers PRJNA931348 and PRJNA931802, respectively.

## Acknowledgments

The sample collection was financially supported by a funding from the Dairy Research Cluster 3 (Dairy Farmers of Canada (Ottawa, ON, Canada) and Agriculture and Agri-Food Canada (Ottawa, ON, Canada)) under the Canadian Agricultural Partnership AgriScience Program and by the Mastitis Network (Saint-Hyacinthe, QC, Canada).

---

**Fig S1.**
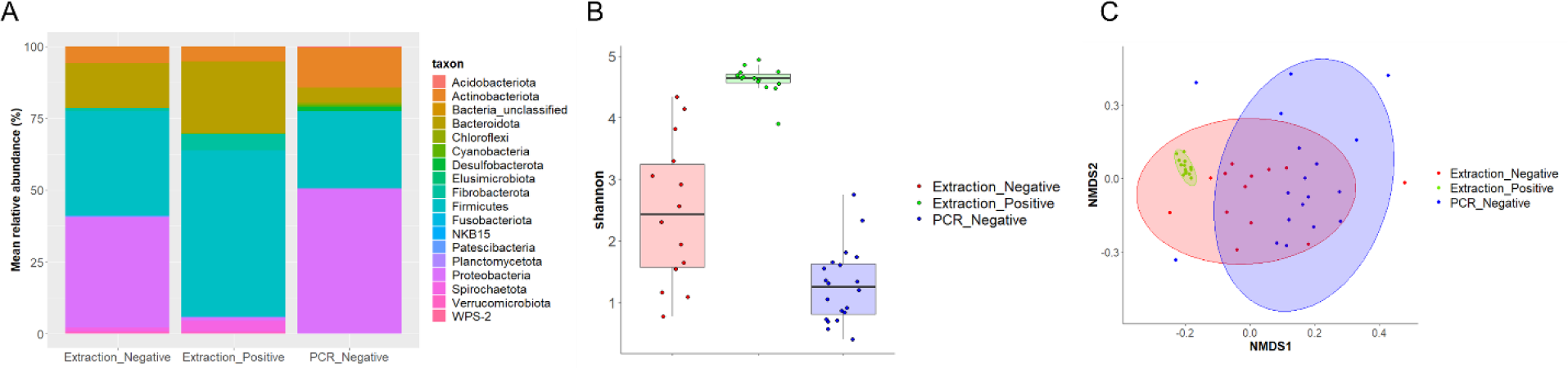
Relative abundance and diversity of the microbiota from the negative and positive controls, and PCR negative control. The DNA extracted from DNA- and RNA-free water was a negative extraction control and a generous donor’s bovine rumen was a positive control for checking contamination from DNA extraction. Positive and negative extraction controls were done for each DNA extraction kit used in the study. In addition, amplification negative controls were also included for the sequencing to ensure no contamination took place at the point of PCR amplification. (A) Each control group had the different relative abundance of phyla. (B) and (C) indicate that Shannon index and Bray-Curtis dissimilarity of each group were significantly different from each other (*p*<0.001). The results indicate there was no contamination during the DNA extraction or amplification.

**Fig S2.**
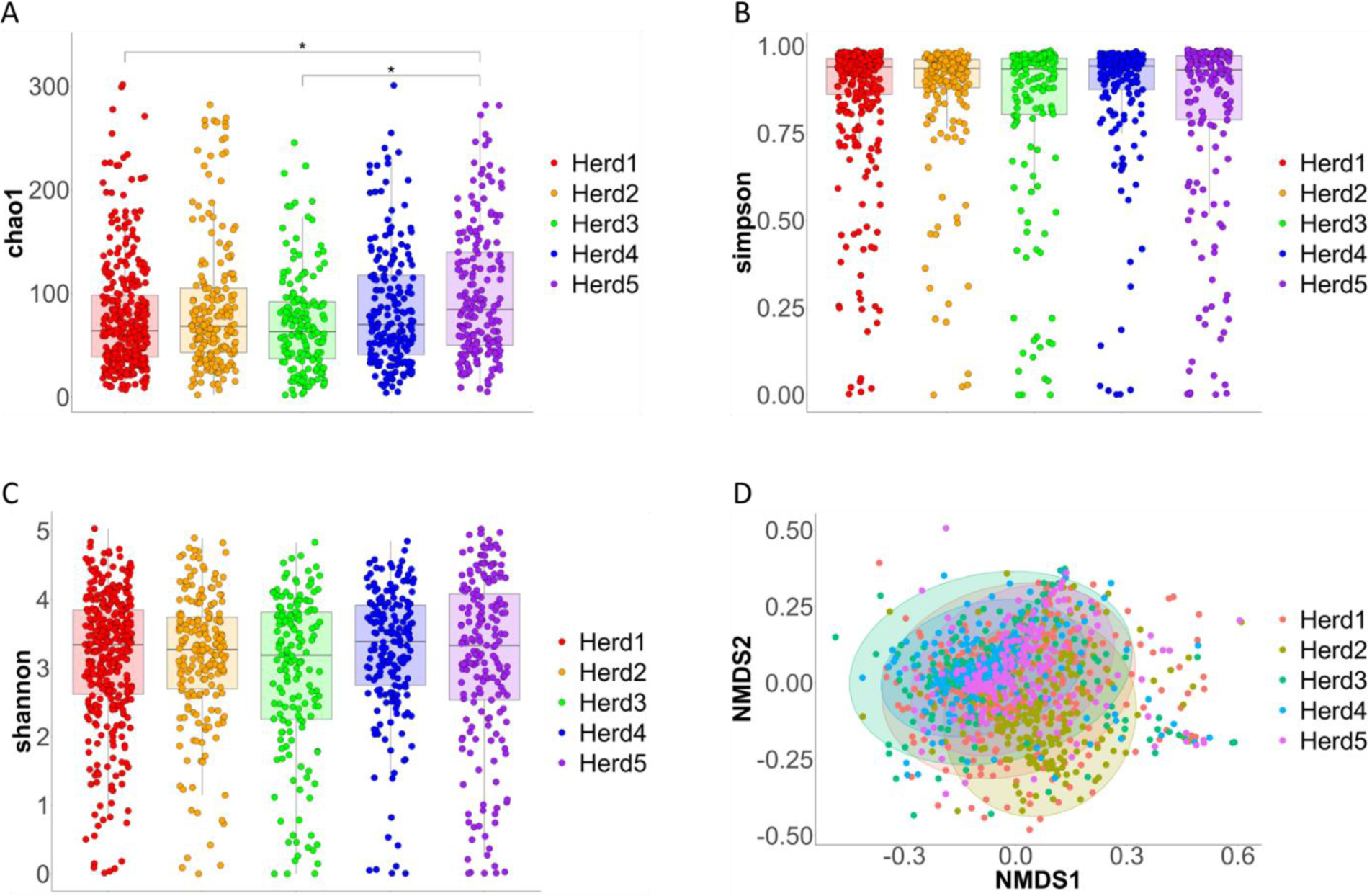
Alpha- and beta-diversities of raw milk microbiota at herd-level. (A), (B) and (C) indicates Chao1, Simpson, Shannon indices of the microbiota from each herd. Significant differences in Chao1 and Simpson indices between herds were identified (One-way ANOVA; *p*<0.001), while Shannon index of each herd was not significantly different (One-way ANOVA; *p*=0.994). (D) Bray-Curtis dissimilarity analysis identified significant difference of beta-diversity between herds (PERMANOVA; *p*<0.001).

**Fig S3.**
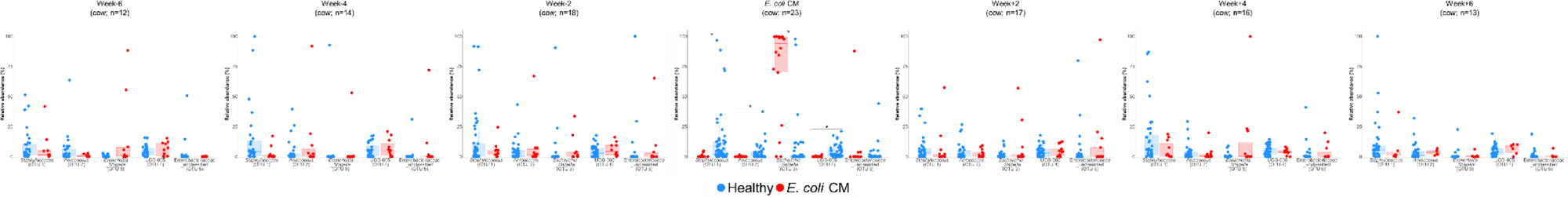
Differential abundance analysis on the five most abundant OTUs in healthy and *E. coli* CM (ECCM) groups using Mann-Whitney U test before, during, and after *E. coli* CM. Mann-Whitney U test adjusted by Benjamini & Hochberg identified significant associations only from *E. coli* CM; OTU0001 (*Staphylococcus*), OTU0002 (*Aerococcus*) and OTU0004 (UCG_005) were significantly associated with healthy samples, and OTU0003 (*Escherichia*_*Shigella*) was significantly associated with ECCM samples (Mann-Whitney U test adjusted by Benjamini & Hochberg; *p* < 0.01)

**Fig S4.**
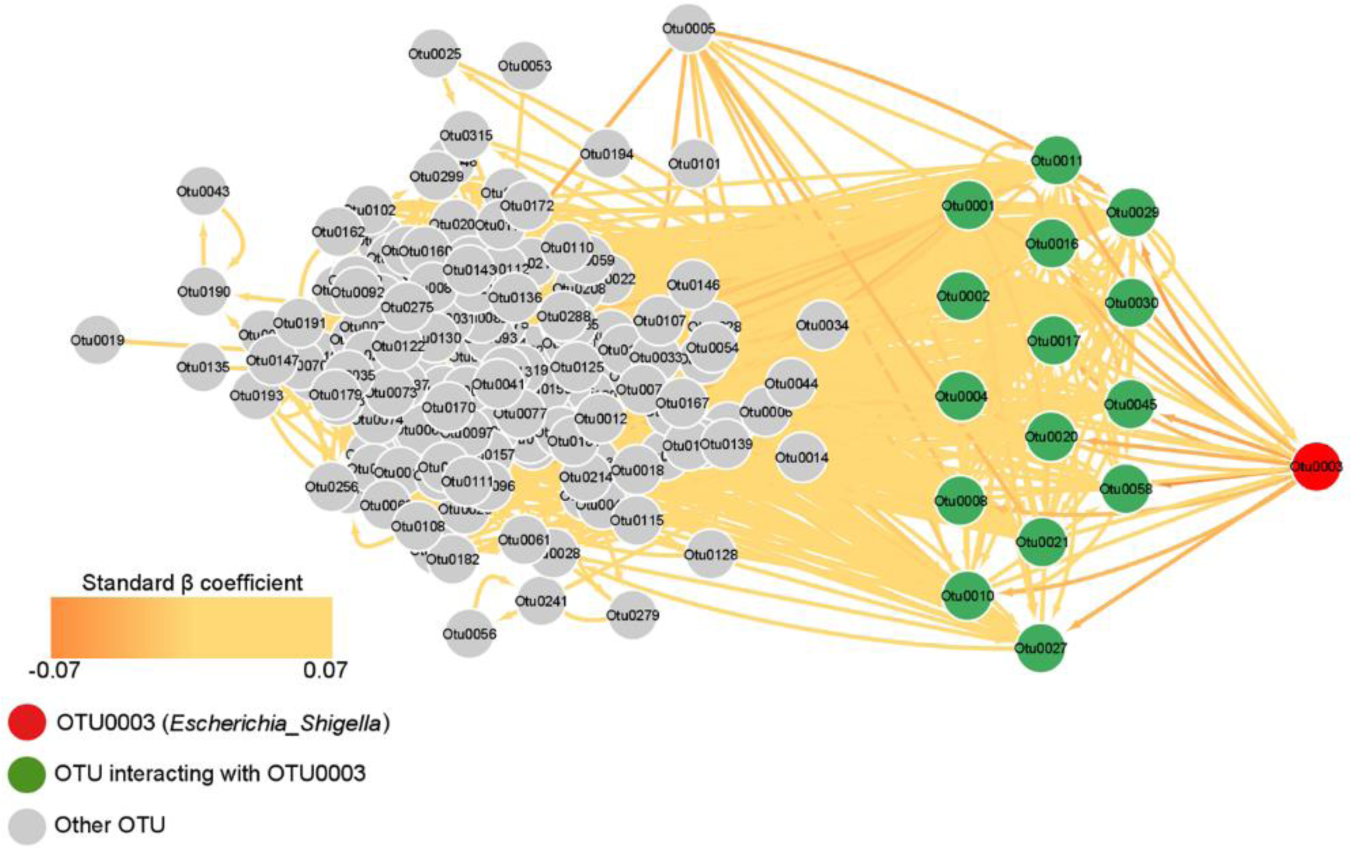
Microbial network of 186 OTUs in the raw milk samples. Combination of classical Spearman correlation and a generalized linear model (GLM) between OTUs created microbial network of the microbiota in 1,172 raw milk samples. Each arrow indicates direction of the relationship based on β calculation and GLM, and the colour of the arrow indicates the strength of either positive or negative relationships based on standard β-coefficient values. In total, 1,795 interactions were detected and 23 of them were interactions between 16 OTUs and OTU0003 (*Escherichia_Shigella*). However, all the interactions were negligible (−0.165 < Spearman ρ < 0.666).

**Fig S5.**
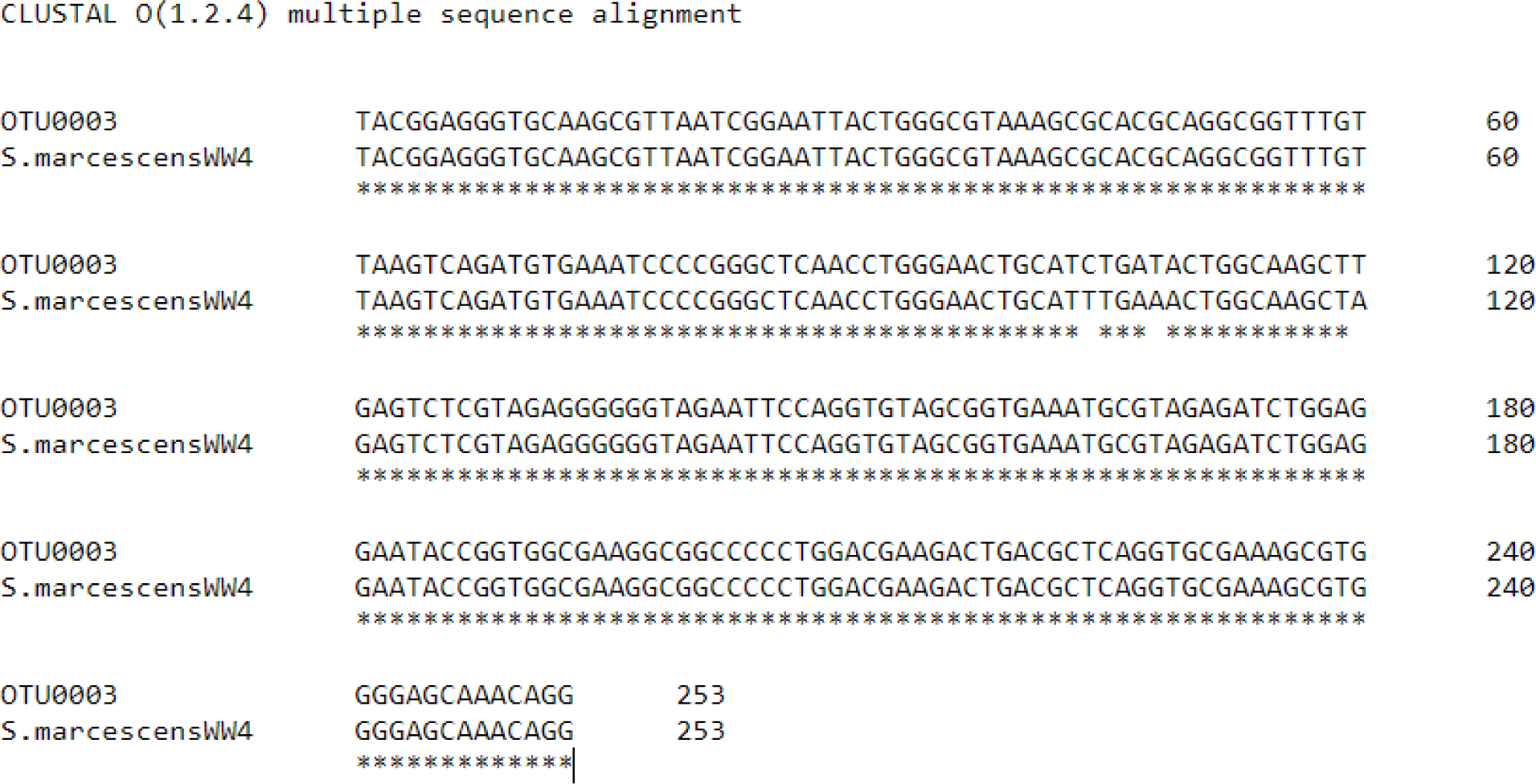
The sequence alignment of V4 region of 16S rRNA gene in OTU0003 and *S. marcescens* WW4. The sequences were aligned using Clustal Omega (v. 1.2.4) (108). The accession number of *S. marcescens* WW4 is PRJNA88659 (109).

